# Coevolution between primates and venomous snakes revealed by α-neurotoxin susceptibility

**DOI:** 10.1101/2021.01.28.428735

**Authors:** Richard J. Harris, K. Anne-Isola Nekaris, Bryan G. Fry

## Abstract

Evidence suggests venomous snakes and primates have evolved certain traits in response to a coevolutionary arms-race. In both clades, evolved traits include an increase in brain size and enhanced vision. Lineage specific traits include in primates an inherent fear of snakes, while cobras have evolved defensive toxins, hooding, aposematism and venom spitting. To strengthen the claims of coevolution between venomous snakes and primates, more evidence of coevolved traits is needed to highlight the importance of this arms-race. We report a significantly reduced susceptibility of snake venom α-neurotoxins toward the α-1 nicotinic acetylcholine receptor orthosteric site within the catarrhine primates. This trait is particularly amplified within the clade Homininae. This relationship is supported by post-synaptic neurotoxic symptoms of envenoming relative to prey species being much lower humans due to weak binding of α-neurotoxins to human nicotinic receptors. Catarrhines are sympatric with many species of large, diurnal, neurotoxically venomous snakes and as such are likely to have had a long history of interaction with them. Conversely, the Lemuriformes and Platyrrhini are highly susceptible to binding of α-neurotoxins, which is consistent with them occupying geographical locations either devoid of venomous snakes or areas with neurotoxic snakes that are small, fossorial, and nocturnal. These data are consistent with the snake detection theory in that they follow a similar pattern of evolved traits within specific primate clades that are sympatric with venomous snakes. These results add new strong evidence in support of snakes and primates coevolving through arms-races that shaped selection pressures for both lineages.

**Significance Statement:** We have discovered a pattern of primate susceptibility towards α-neurotoxins that supports the theory of a coevolutionary arms-race between venomous snakes and primates.

## Introduction

There is growing evidence that coevolutionary interactions between primates and snakes have shaped primate neurobiology, psychology and physiology; this is known as the snake detection theory of primate evolution(1). The evidence proposed in this theory suggests that constricting snakes were the first predators of crown-group placental mammals including early primates and that the ‘fear module’(2), visual specialisation (e.g. object recognition) and brain expansion of primates all evolved in response to early constricting snakes. A further visual system refinement (e.g. trichromacy) later evolved in anthropoids, including Catarrhini (African/Asian monkeys, apes and humans) and Platyrrhini (American monkeys), as a response to the evolution of venomous snakes such as Elapidae and Viperidae(1). Further, a greater visual bias leading to greater detection of snakes than to other animals/objects has been found in catarrhines including *Homo sapiens(3–7).* These adaptations differ between major lineages of primates and coincide with the evolutionary co-existence with venomous snakes. For example, primates of the suborder Strepsirrhini (Lorisiformes, Lemuriformes) have visual systems characterised by mono- or dichromacy, whereas all catarrhines and some platyrrhines are characterised by trichromacy(1). Lemuriformes colonised Madagascar between 50-65 Ma(8). Thus they likely were not sympatric with vipers (~50 MA)(9), cryptic, highly camouflaged, ambush feeders that would have had little interaction with primates in any case. Most importantly, regardless of divergence date, lemurs were not likely to have been sympatric with the neurotoxic elapids (~37 Ma)(10). Madagascar has remained devoid of venomous snakes, thus there has never been a selection pressure on this group to evolve survival adaptations against venomous snakes. The platyrrhines on the other hand have a highly variable visual systems, with both dichromacy and trichromacy prevalent within the group(1). Platyrrhines occupied South America ~35 Ma(11), which is long before the arrival of venomous snakes between 3-25 Ma, based on both molecular data and the formation of the Panamanian land bridge(9, 12–14). Thus, platyrrhines have had an interrupted evolutionary time scale with venomous snakes, with the recently sympatric venomous snakes being mostly terrestrial, and with neurotoxic species in particular being fossorial and nocturnal. Furthermore, selection pressures of terrestrial snakes to these largely arboreal primates would be negligible.

Conversely, there is evidence that venomous elapids have evolved some adaptations in response to primates. Spitting in cobras has evolved on three convergent occasions (twice within *Naja* and again *Hemachatus)* is a defensive trait that causes intense ocular pain and inflammation, which might have evolved in response to unique behavioural adaptations by anthropoids, particularly early hominines(15). Predation events by natural predators (e.g. birds and other reptiles) of spitting cobras often occur, thus it seems that spitting is not an effective method against most predators(15). Intriguingly, the emergence of spitting in the *Hemachatus* (<17 mya), the African spitting *Naja* (~6.7 mya), and the Asian spitting *Naja* (~2.5 mya)(10), all coincided with the diverse emergence of terrestrial dwelling apes, including early hominids(16, 17), all of which occurred in habitats also frequented by largely terrestrial spitting cobras(18). These hominids showed a myriad of features indicating a tendency towards upright bipedal movement and increased precision and opposability of the hands and thumbs. It has been inferred that the last common ancestors of chimpanzees and humans already used wood and stone tools and had the ability to hunt vertebrates(17). Furthermore, terrestrial dwelling apes, through their locomotor style, foraging habits, and groups movements, are more likely than other terrestrial taxa to come into frequent contact with venomous snakes(19). The emergence of Asian spitting cobras at ~2.5 mya(10) coincides with the appearance of *Homo erectus* (~1.8 mya), the latter of which is characterised by advancements in stone tool technology(20). Primates in general show overt reactions to snakes, including snake-specific alarm calls, and may mob venomous snakes with improvised tools and projectiles(21–23). Thus, the use of projectile weaponry in primates, particularly in anthropoids that stand upright when using such weapons, would provide a selection pressure on snakes to evolve a ranged defense such as spitting venom at the eyes of upright primates. Additionally, the evolution of injury inducing defensive cytotoxins is also correlated with that of defensive hooding displays and aposematic marking in African and Asian cobras(24). The evolution of defensive cytotoxins also preceding that of spitting suggests that a high selection pressure might have been exerted upon cobras for defense against primates before the emergence of spitting when hominins evolved bipedalism. Hence, there is some fundamental evidence to consider that reciprocal coevolution occurred between primates and snakes, particularly venomous snakes, driving some novel traits.

Predator-prey relationships between venomous snakes and non-primate mammals is a widely researched area, particularly regarding resistance mechanisms evolved by mammals toward the venom of snakes (25–30). A prominently evolved resistance mechanism is seen at the orthosteric site (acetylcholine binding region) of the α-1 nicotinic acetylcholine receptor (nAChR) by which specific amino acid residues reduce the binding of α-neurotoxins (such as three-finger toxins; 3FTxs) found within some snakes venoms, particularly that of Elapidae(31, 32). The *H. sapiens* α-1 nAChR is much less susceptible to binding of α-neurotoxins than other organisms(33–36). This might also be a reason why post-synaptic neurotoxicity of snakebites from a large proportion of elapid species in humans is rarely an observed clinical symptom, but prey items are rapidly paralysed. Despite this, no studies have attempted to investigate further this reduced susceptibility in *H. sapiens* and how α-neurotoxin binding might affect a wide range of primate species, given the surmounting evidence in support of their coevolution with venomous snakes.

Investigations into the publicly available α-1 nAChR sequences of a range of primate species revealed that the orthosteric site of *H. sapiens* is conserved across the Homininae (chimpanzees, gorillas, humans), whilst other primates such as Cercopithecidae (African/Asian monkeys), Galagidae (bushbabies), Lemuriformes (lemurs), Platyrrhini and Ponginae (orangutans) all have clade-specific orthosteric sequences (Table S1). Given these differences between primate clades and the theory that primates and snakes might have coevolved certain traits through reciprocal antagonism, we tested if there was any differences in α-neurotoxin binding between primate lineages, particularly given evidence that *H. sapiens* nAChRs are less susceptible to α-neurotoxins(33–36). To assess these differences, we tested five venoms that are rich in α-neurotoxins (3FTxs) from species of cobra *(Naja* spp.) that both inhabit Africa (*N. haje, N. mossambica* and *N. nubiae)* and Asia (*N. kaouthia* and *N. siamensis).* These species were further categorised as spitting (*N. mossmabica, N. nubiae* and *N. siamenis)* and non-spitting *(N. haje* and *N. kaouthia)* species. We tested the binding of these venoms upon mimotopes that represent the orthosteric site of nAChRs from different primate clades (Table S1). We utilised a biolayer interferometry assay (BLI), that has been previously validated to assess the binding of α-neurotoxins to taxa specific orthosteric mimotopes(33, 37–39) to determine if coevolutionary interactions between cobras and primates might have evolved some resistance elements toward α-neurotoxins.

## Results and Discussion

The results indicate a consistent binding pattern across all *Naja* venoms tested toward the primate lineage mimotopes (Fig 1–5). The non-spitting species seem to have a much lower binding overall than the spitting, which would suggest lower proportions of 3FTxs within their venom(37). The difference observed in binding of the crude venoms is a consequence of the toxin interactions with the different biochemical properties of each amino acid present within the sequences(35, 40–42). The binding patterns across all venoms tested showed a significantly reduced susceptibility of α-neurotoxins across the Homininae (Fig 1–5). These data additionally support previous observations of weakly binding α-neurotoxins toward human nAChRs(33–36). Further, the venoms were also bound relatively weakly to the Cercopithecidae and Ponginae mimotopes (Fig 1–5). These results are interesting since Cercophitecidae, Homininae and Ponginae make up the clade Catarrhini that remained and evolved in Africa and later spread out through Asia similarly to cobras, suggesting that coevolutionary selection pressures were being maintained across this distribution. The Lorisiformes and Tarsiiformes representatives also have relatively low susceptibility, and both of these clades also occupy Africa and Asia respectively, regions in which there are an abundance of α-neurotoxic snake species. Both these groups also have particularly high orbital convergence, which has been associated with their co-evolution with snakes(1). Within the Lorisidae, the Sunda slow loris *(Nycticebus coucang)* contains the N-glycosylation motif(43–46) (Table S1) that has been associated with α-neurotoxin resistance, which is consistent with them being documented as eating venomous snakes. Lorises are also venomous themselves (47, 48), but given the extreme cytotoxicity of their venom and lack of neurotoxic symptoms, the N-glycosylation it is unlikely to have evolved for autoresistance.

**Fig. 1.**
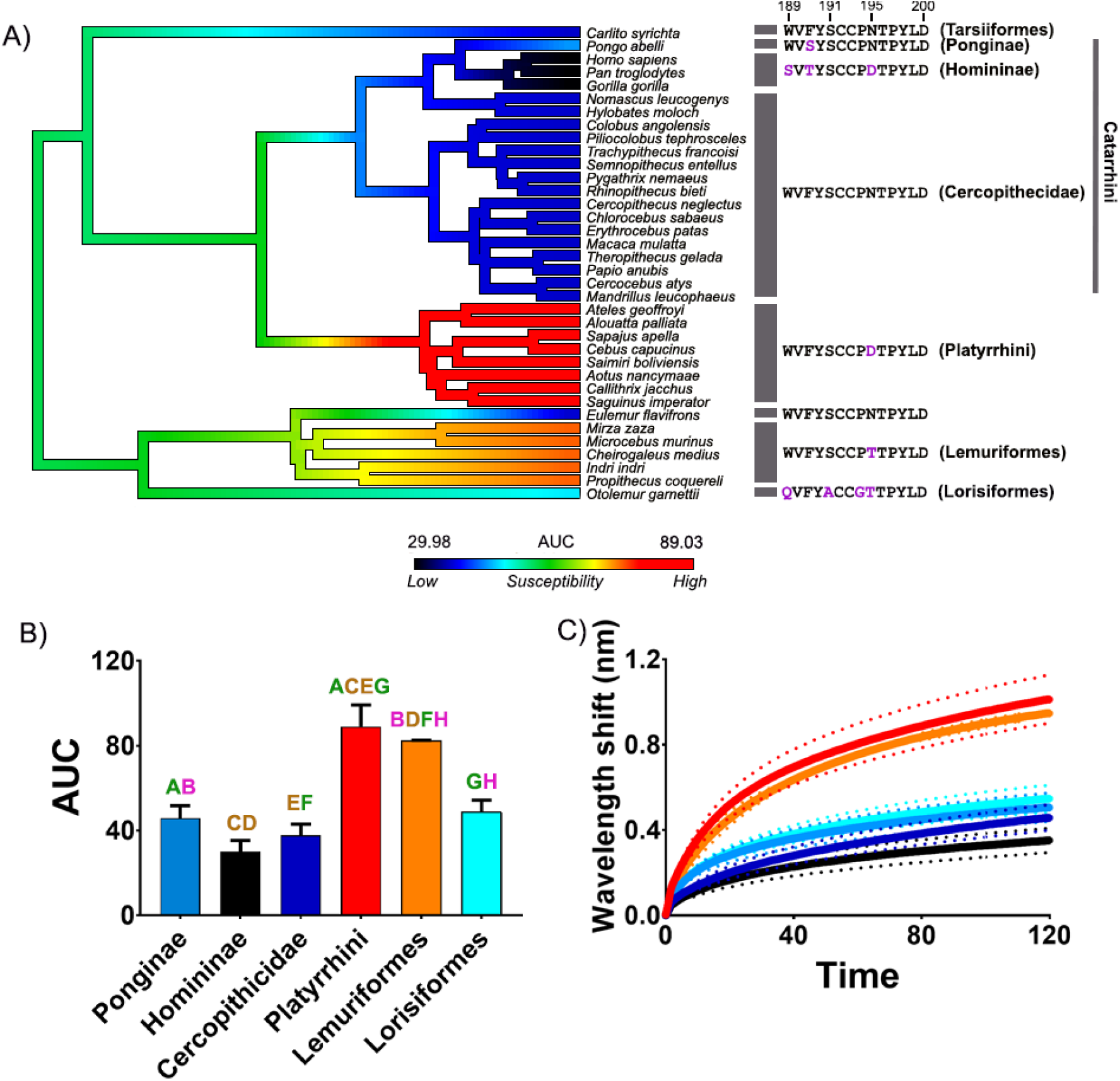
The effects of venom from the African cobra *Naja mossambica* against the nAChR orthosteric site mimotopes from seven clades of primates. A) Ancestral state reconstruction of the area under the curve (AUC) values of the binding of *N. mossambica* against the primate mimotopes. B) Bar graphs represent the mean area under the curve (AUC) values of the adjacent curve graphs. C) Curve graphs show the mean wavelength shift (nm) in light with binding of venoms over a 120 second association phase. The venom was tested in triplicate (n=3). Error bars on all graphs represent the SEM. AUC values were statistically analysed using a one-way ANOVA with a Tukey’s comparisons multiple comparisons test comparing to the native mimotope. Statistical significance is indicated by matching letters with the colours of letter indicating the level of significance; brown p<0.001, green p<0.01, pink p<0.05. All raw data and statistical analyses outputs can be found in Supplementary data S1.

**Fig. 2.**
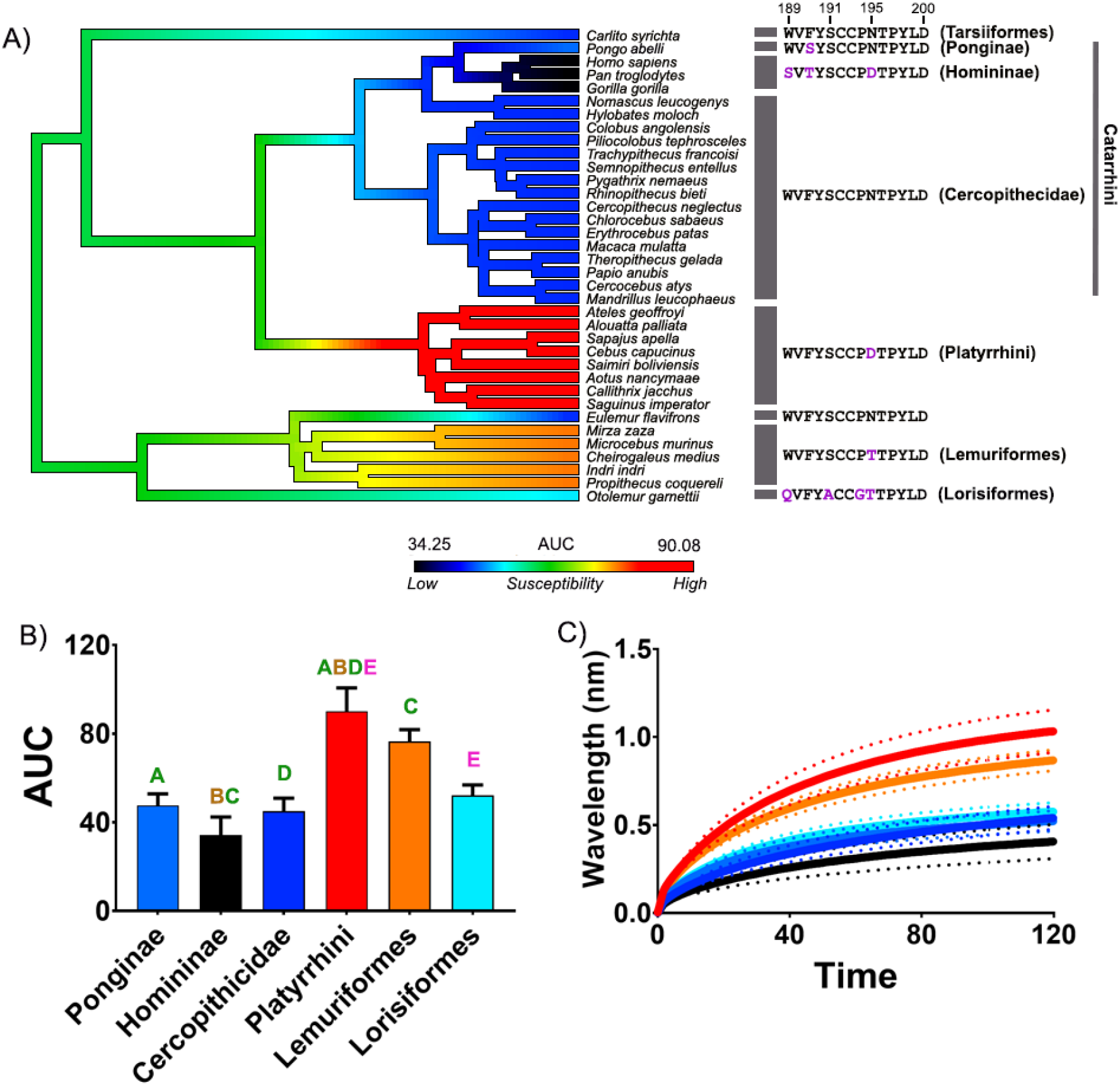
The effects of venom from the African cobra *Naja nubiae* against the nAChR orthosteric site mimotopes from seven clades of primates. A) Ancestral state reconstruction of the area under the curve (AUC) values of the binding of *N. nubiae* against the primate mimotopes. B) Bar graphs represent the mean area under the curve (AUC) values of the adjacent curve graphs. C) Curve graphs show the mean wavelength shift (nm) in light with binding of venoms over a 120 second association phase. The venom was tested in triplicate (n=3). Error bars on all graphs represent the SEM. AUC values were statistically analysed using a one-way ANOVA with a Tukey’s comparisons multiple comparisons test comparing to the native mimotope. A statistical significance is indicated by matching letters with the colours of letter indicating the level of significance; brown p<0.001, green p<0.01, pink p<0.05. All raw data and statistical analyses outputs can be found in Supplementary data S1.

**Fig. 3.**
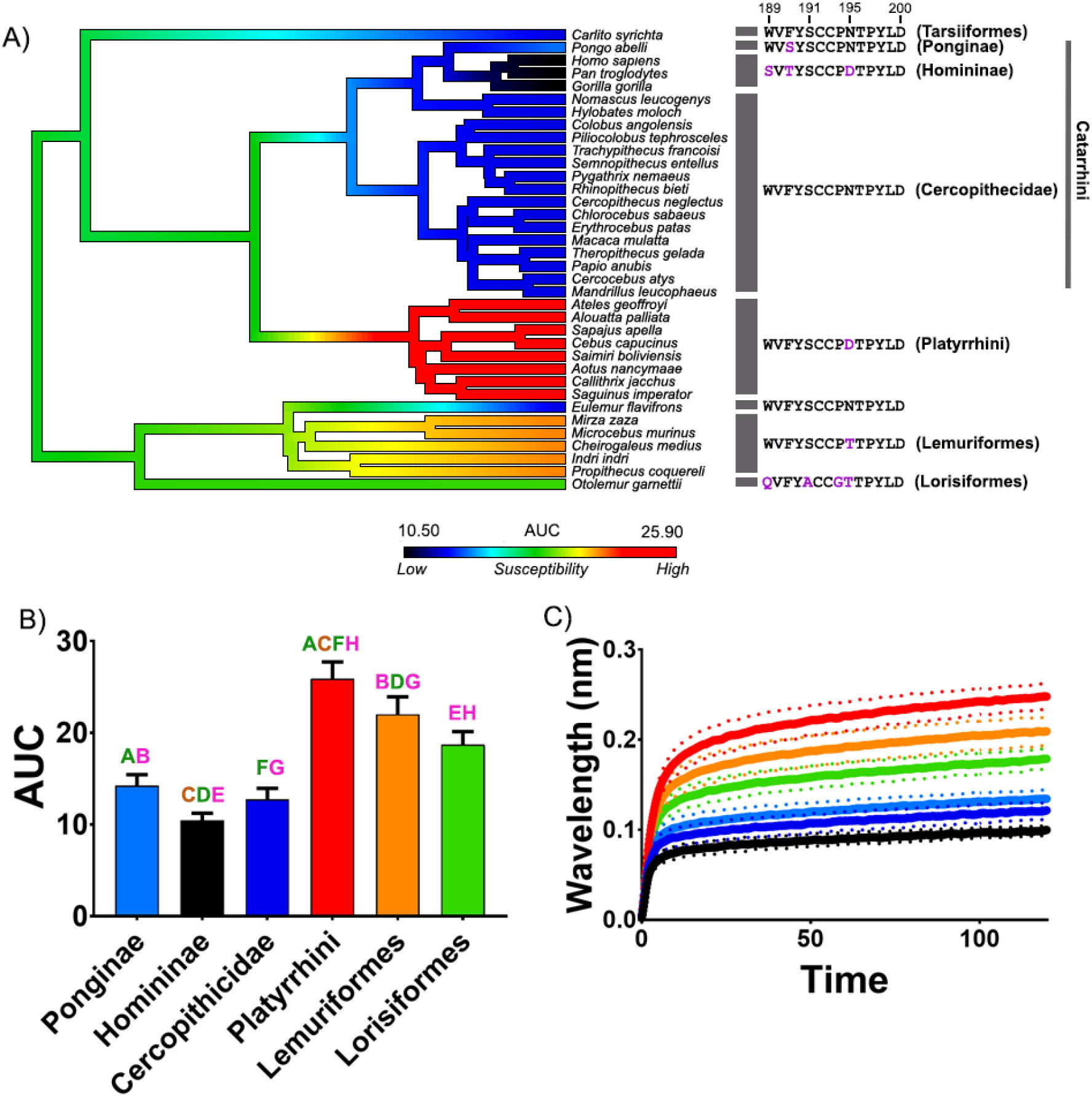
The effects of venom from the African cobra *Naja haje* against the nAChR orthosteric site mimotopes from seven clades of primates. A) Ancestral state reconstruction of the area under the curve (AUC) values of the binding of *N. haje* against the primate mimotopes. B) Bar graphs represent the mean area under the curve (AUC) values of the adjacent curve graphs. C) Curve graphs show the mean wavelength shift (nm) in light with binding of venoms over a 120 second association phase. The venom was tested in triplicate (n=3). Error bars on all graphs represent the SEM. AUC values were statistically analysed using a one-way ANOVA with a Tukey’s comparisons multiple comparisons test comparing to the native mimotope. A statistical significance is indicated by matching letters with the colours of letter indicating the level of significance; brown p<0.0001, green p<0.001, pink p<0.05. All raw data and statistical analyses outputs can be found in Supplementary data S1.

**Fig. 4.**
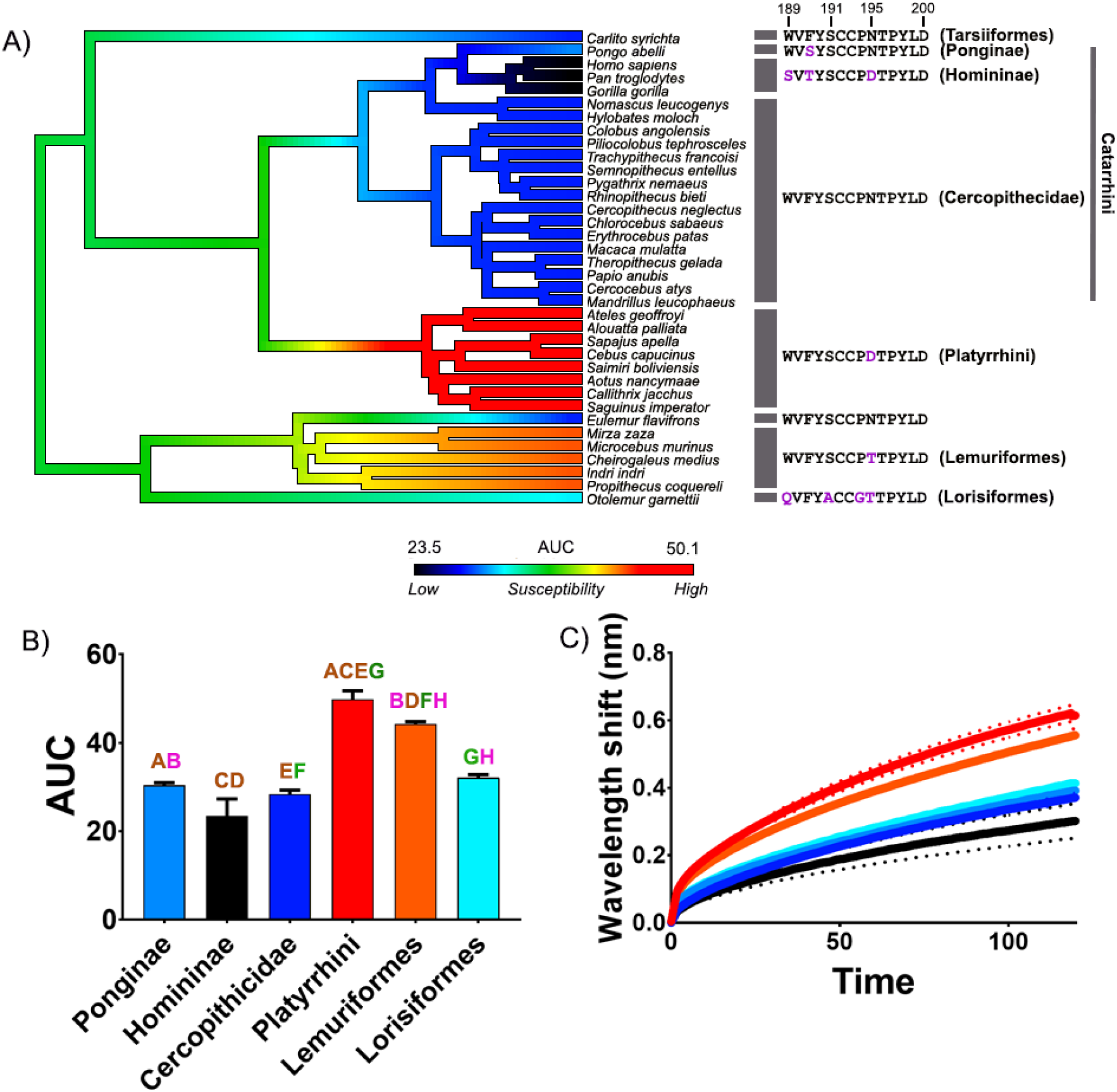
The effects of venom from the Asian cobra *Naja siamensis* against the nAChR orthosteric site mimotopes from seven clades of primates. A) Ancestral state reconstruction of the area under the curve (AUC) values of the binding of *N. siamensis* against the primate mimotopes. B) Bar graphs represent the mean area under the curve (AUC) values of the adjacent curve graphs. C) Curve graphs show the mean wavelength shift (nm) in light with binding of venoms over a 120 second association phase. The venom was tested in triplicate (n=3). Error bars on all graphs represent the SEM. AUC values were statistically analysed using a one-way ANOVA with a Tukey’s comparisons multiple comparisons test comparing to the native mimotope. A statistical significance is indicated by matching letters with the colours of letter indicating the level of significance; brown p<0.0001, green p<0.001, pink p<0.01. All raw data and statistical analyses outputs can be found in Supplementary data S1

**Fig. 5.**
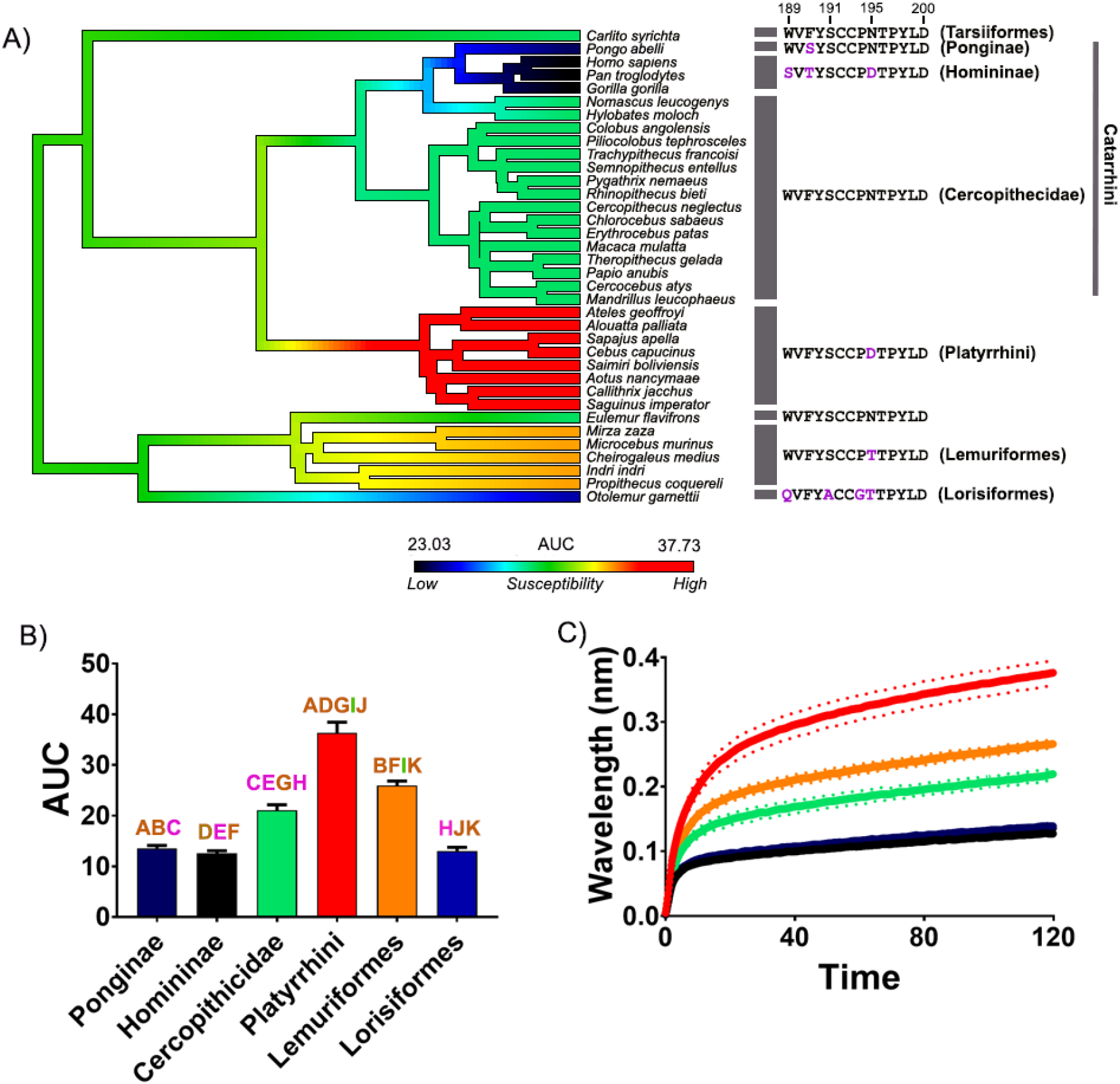
The effects of venom from the Asian cobra *Naja kaouthia* against the nAChR orthosteric site mimotopes from seven clades of primates. A) Ancestral state reconstruction of the area under the curve (AUC) values of the binding of *N. kaouthia* against the primate mimotopes. B) Bar graphs represent the mean area under the curve (AUC) values of the adjacent curve graphs. C) Curve graphs show the mean wavelength shift (nm) in light with binding of venoms over a 120 second association phase. The venom was tested in triplicate (n=3). Error bars on all graphs represent the SEM. AUC values were statistically analysed using a one-way ANOVA with a Tukey’s comparisons multiple comparisons test comparing to the native mimotope. A statistical significance is indicated by matching letters with the colours of letter indicating the level of significance; brown p<0.0001, green p<0.001, pink p<0.05. All raw data and statistical analyses outputs can be found in Supplementary data S1.

Conversely, the Lemuriformes (with the exception of *Eulemur flavifrons)* and Platyrrhini clades were the most susceptible to binding by the venoms (Fig 1–5). Both these clades were geographically separated from the African and Asian primate clades and thus also from large venomous snake species that would potentially provide selection pressures. The Lorisiformes did not migrate to Madagascar and persisted under higher selection pressures from α-neurotoxic elapids in Africa and Asia.

These data suggest that there is a selection pressure that has led to African and Asian primates becoming less susceptible toward cobra α-neurotoxins than other primate clades that are geographically separated. The emerging patterns seem consistent with the snake detection theory(1) in that the primate clades that evolved in Africa and/or remained sympatric to venomous snakes seem to have evolved mechanisms to cope with certain selective pressures. Catarrhini, Lorisiformes and Tarsiiformes clades have evolved some reduced susceptibility toward α-neurotoxins whereas the Malagasy Lemuriformes have never co-existed with venomous snakes and the Platyrrhini occupy Central and South America where the elapids are small, nocturnal and fossorial, and thus pose little threat.

From an evolutionary perspective it would seem that the reduced susceptibility across African and Asian primates is likely a basal trait and the increase in susceptibility has independently evolved with the major lineage splits of Lemuriformes and Platyrrhini (Fig 1–5). Since the basal trait seems to be a reduced susceptibility it is likely that there is an evolutionary trade-off disadvantage to fitness with evolving specific residue changes that lowers α-neurotoxin binding such as a reduced binding of the endogenous acetylcholine. This is possibly why both Lemuriformes and Platyrrhini have evolved substitutions that increase α-neurotoxin susceptibility as there is no longer a sufficient selection pressure to keep the ancestral state. In the case of *E. flavifrons,* it is uncertain why this species did not evolve a similar sequence to the rest of the Lemuriformes, conspicuously some dietary items of *Eulemur,* such as mushrooms and millipedes, contain high levels of neurotoxins (49). Future work should investigate if the toxins from these sources target nAChRs (e.g. high concentrations of nicotine) and provide enough of a selection to maintain the lower susceptibility toward nAChR targeting toxins.

It is important to note that primates are a primarily arboreal order, and even in the trees, snakes are a major antagonist to primates, resulting in a strong fear module against snakes(1). The period in which spitting in cobras evolved occurred during a major climatic shift linked to reduction in continuous forests(17). During this time, diurnal primates in particular would have needed to descend to the ground, and a number of species adapted to be fully terrestrial, increasing their chance of encountering and being defensively bitten by venomous snakes. Considering that snakes are not prey to frugivorous diurnal primates, but are a danger if startled, it is argued that snakes placed a tremendous selection pressure on primates, including their parvocellular pathway, providing early visual detection of snakes, and allowing subsequent fight or flight responses(1).

The greater reduction in susceptibility within Homininae coincides not only with obligate bipedalism and reduced time in trees, but also with the reduction of continuous forests requiring movements into more open habitats. Foraging hominins were likely to come in contact much more often with terrestrial, diurnally active neurotoxic elapid snakes such as cobras, which would defend themselves against perceived primate predators using their venom(22). The likely reduced susceptibility to α-neurotoxins from cobras might have driven cobras to develop a different kind of defense against hominins in the form of cytotoxins and spitting(15, 24). An evolved form of partial resistance meant that cobra venom was not an effective method to reduce hominin interactions thus cytotoxins and spitting might have evolved in response as a deterrent. Since it has been proposed that spitting cobra clades are a major co-evolutionary counterpart to primates, this might explain the difference we observed in binding between spitting and nonspitting cobras. A lower binding of α-neurotoxins in the venom of non-spitting than spitting cobras (likely due to the proportion of α-neurotoxins within the venom), might be that the reduced susceptibility across African and Asian primates has meant that spitters have had to evolve a high proportion of α-neurotoxins to overcome this reduced susceptibility. Yet this proposed hypothesis needs careful consideration and investigation.

Further, although predation of primates from snakes does occur(50), albeit rarely, it is more likely that defensive bites from venomous snakes is the driving selection pressure. Defensive scenarios between venomous snakes and primates have been observed, for example, two female chacma baboons *(Papio hamadryas ursinus)* were bitten and died from a defensive cape cobra *(Naja nivea)* bite, whilst another individual, who recovered from a bite, was spat in the eyes by the venom(51).

Thus, given that commonly encountered neurotoxic species of snakes exert a strong selection pressure in their defensive envenomations, why does there only seems to be a reduced susceptibility rather than full resistance? Firstly, our aforementioned proposition that there might be an evolutionary trade-off disadvantage to evolving full resistance at the nAChR orthosteric site whereby the endogenous neurotransmitter acetylcholine does not bind as efficiently due to resistance mechanisms employed. Therefore, evolving reduced susceptibility might allow for a balance between partial resistance and efficient acetylcholine binding. This is supported by previous work revealing that vipers which are prey to neurotoxic snakes often display resistance, but this resistance is secondarily lost in populations which have radiated out into geographic areas devoid of neurotoxic snakes(46). Secondly, since primates are social animals(52, 53) it is possible that reduced susceptibility might have been successful enough for kin selection(54–56), to maintain the partial resistance trait reciprocally. Having a reduced susceptibility would allow the envenomed individual to fight against the snake for longer than more susceptible primates and also warn other members of the group that a venomous snake has entered the territory. Warning calls towards snakes occur within both nocturnal and diurnal primate groups(57, 58), thus being able to marginally extend an individual’s survival time to warn the group, allowing the safety of the relatives and, ultimately, the successful reproduction of the group.

In summary we have revealed a reduced susceptibility across African and Asian primates, particularly within the catarrhines, towards snake venom α-neurotoxins that target nicotinic acetylcholine receptors. Homininae is the least susceptible when tested across African and Asian spitting and non-spitting cobras. This finding is consistent with clinical symptoms of neurotoxicity being rare in envenomations by α-neurotoxic snakes. Further to this, the Lemuriformes and Platyrrhini have the highest susceptibility of primates tested toward α-neurotoxic venoms. This finding is consistent with the evolutionary biogeography of these lineages, which have geographically radiated to Madagascar and South America respectively, and evolved in the absence of large neurotoxic snake species. Our data also further supports one of the hypothesis proposed within the snake detection theory of primate evolution, whereby venomous snakes and primates share a history of coevolution thorough their continuous co-existence across Africa and asia(1). The evidence in support of the coevolution of primates and snakes seems to be increasing, yet despite this there are still fundamental gaps in knowledge both regarding the ecology of both snakes and primates and how their shared coevolutionary history might have brought about some of their most distinguishing traits.

## Materials and Methods

### Venom collection and preparation

Venoms were sourced from individual adult snakes (captive and wild-caught) from a long-term cryogenic collection. All venom samples were lyophilised and reconstituted in double deionised water (ddH_2_O), and centrifuged (4 °C, 10 min at 14,000 relative centrifugal force (RCF)). The supernatant was made into a working stock (1 mg/mL) in 50% glycerol at −20 °C. The concentrations of experimental stocks were determined using a NanoDrop 2000 UV-Vis Spectrophotometer (Thermo Fisher, Sydney, Australia) at an absorbance wavelength of 280 nm.

### Mimotope production and preparation

Following methods from a previously developed assay (37, 38), a 13-14 amino acid mimotope of the vertebrate α-1 nAChR orthosteric site was developed by GenicBio Ltd. (Shanghai, China) designed upon specification. Mimotopes from lineage representatives were as follows; *Homo sapiens* (G5E9G9), *Otolemur garnettii* (H0WHF2), *Pongo abelli* (H2P7W2), lemur (P04756), Cercopithecidae (P25108) and platyrrhini *Nomascus leucogenys* (G1T5.3). The full list is available as Supplementary Dataset 1.

The C-C of the native mimotope is replaced during peptide synthesis with S-S to avoid uncontrolled postsynthetic thiol oxidation. The C-C bond in the nAChR binding region does not participate directly in analyte-ligand binding(42, 59, 60), thus replacement to S-S is not expected to have any effect on the analyte-ligand complex formation. However, the presence of the C-C bridge is key in the conformation of the interaction site of whole receptors(61). As such, we suggest direct comparisons of kinetics data, such as Ka or KD, between nAChR mimotopes and whole receptor testing should be avoided, or at least approached with caution. Mimotopes were further synthesised to a biotin linker bound to two aminohexanoic acid (Ahx) spacers, forming a 30 Å linker.

Mimotope dried stocks were solubilised in 100% dimethyl sulfoxide (DMSO) and diluted in ddH_2_O at 1:10 dilution to obtain a stock concentration of 50 μg/mL. Stocks were stored at −80 °C until required.

### Biolayer Interferometry (BLI)

Full details of the developed assay, including all methodology and data analysis, can be found in the validated protocol(38) and further data using this protocol(33, 37). In brief, the BLI assay was performed on the Octet HTX system (ForteBio, Fremont, CA, USA). Venom samples were diluted 1:20, making a final concentration of 50 μg/mL per well. Mimotope aliquots were diluted 1:50, with a final concentration of 1 μg/mL per well. The assay running buffer was 1X DPBS with 0.1% BSA and 0.05% Tween-20. Preceding experimentation, Streptavidin biosensor were hydrated in the running buffer for 30-60 min, whilst on a shaker at 2.0 revolutions per minute (RPM). The dissociation of analytes occurred using a standard acidic glycine buffer solution (10 mM glycine (pH 1.5-1.7) in ddH_2_O). Raw data are provided in Supplementary files.

### Data processing and analysis

All raw data is available in Supplementary Dataset 2. All data obtained from BLI on Octet HTX system (ForteBio) were processed in exact accordance to the validation of the previously validated assay(38). The association step data were obtained and imported into Prism8.0 software (GraphPad Software Inc., La Jolla, CA, USA) where area under the curve (AUC) and one-way ANOVA with a Tukeys’s multiple comparisons analyses were conducted and graphs produced.

## Notes

### Competing Interest Statement

The authors have declared no competing interest.

